# On the relation between input and output distributions of scRNA-seq experiments

**DOI:** 10.1101/2021.10.08.463628

**Authors:** Daniel Schwabe, Martin Falcke

**Affiliations:** Mathematical Cell Physiology, Max Delbrück Center for Molecular Medicine in the Helmholtz Association, Robert-Rössle-Str. 10, 13125 Berlin, Germany; Department of Physics, Humboldt University Berlin, Newtonstr. 15, 12489 Berlin, Germany

## Abstract

**Motivation:** Single-cell RNA sequencing determines RNA copy numbers per cell for a given gene. However, technical noise poses the question how observed distributions (output) are connected to their cellular distributions (input).

**Results:** We model a single-cell RNA sequencing setup consisting of PCR amplification and sequencing, and derive probability distribution functions for the output distribution given an input distribution. We provide copy number distributions arising from single transcripts during PCR amplification with exact expressions for mean and variance. We prove that the coefficient of variation of the output of sequencing is always larger than that of the input distribution. Experimental data reveals the variance and mean of the input distribution to obey characteristic relations, which we specifically determine for a HeLa data set. We can calculate as many moments of the input distribution as are known of the output distribution (up to all). This, in principle, completely determines the input from the output distribution.

## Introduction

When exploring cell populations, one might expect that cells of the same type and under the same environmental conditions should have the exact same gene expression profile. However, single-cell data has revealed that gene expression even among clonal cells is heterogeneous. This is called biological noise or cell-to-cell variability. While causes for this effect have been discussed extensively [8, 26, 31], the consequences or intent of it are more obscure [23, 24, 26]. Nevertheless, it appears frequently in data sets such that quantification becomes more and more important.

Due to cell-to-cell variability [35], we face distributions of individual mRNA species across a cell population instead of a single abundance value for all of them. That true distribution of the abundance of a specific gene is the input distribution to single-cell RNA sequencing (scRNA-seq). The scRNA-seq experiment will then introduce significant technical noise [13, 36] and produce an observed or output distribution. The technical and biological noise are convoluted in the output distribution. We have to remove the technical noise from the data in order to estimate cell-to-cell variability. Many data-driven approaches have been established to deal with various aspects of technical noise. Among others, normalization [2, 5, 6, 10, 22], batch correction [6, 11, 34] and data imputation [14, 15, 20, 37] are now often part of scRNA-seq data analysis in an attempt to remove technical noise.

In contrast to such a data-driven approach, we model a simplified version of a scRNA-seq experiment and propose probability distributions for the data at each step. This results in an analytical formula for the output distribution for a given input distribution. Analysing this analytic expression reveals the moments of the input distribution, which is a first step towards quantifying cell-to-cell variability.

We simplify scRNA-seq experiments to the following essential steps:

1. Starting point is a cell population of *n* cells each containing *m* genes.
2. The mRNA is extracted and tagged with a cell barcode and a unique molecular identifier (UMI) sequence.
3. The mRNA material is amplified with *l* cycles of PCR.
4. The mRNA library is sequenced and reads containing the same UMI are collapsed.

### The PCR distribution

Since each individual transcript is tagged by a cell barcode and a UMI, it can be uniquely identified amongst all other transcripts. The likelihood to produce the same cell barcode or UMI twice is small enough to justify ignoring these occurrences in our setting.

PCR is a method to amplify DNA material. At each cycle, every molecule has a chance *p* to be copied once. *p* is called the PCR efficiency. In an ideal scenario, *p* would be equal to 1. In reality, efficiency values vary greatly and the importance of having perfect efficiency depends entirely on the research question. Typical estimates put PCR efficiency between 0.9 and 1 [32]. While the PCR efficiency can be different from cycle to cycle [4], we simplify our estimates and assume that the PCR efficiency *p* is constant across all cycles.

We now want to know how many PCR copies each unique initial molecule produces after *l* cycles. Let *X*_*l*_ be the random variable describing the number of copies after *l* cycles with *X*_0_ = 1. The increase of copy numbers from *X*_*l*−1_ to *X*_*l*_ during cycle *l* follows a binomial distribution since each existing copy has chance *p* to be copied again. This means (*X*_*l*_ −*X*_*l*−1_|*X*_*l*−1_ = *n*) ∼ Binom(*n, p*). Therefore, we can recursively derive an analytical formula for the probability distribution function (pdf) ℙ[*X*_*l*_ = *k*] to obtain *k* copies after *l* cycles. This yields

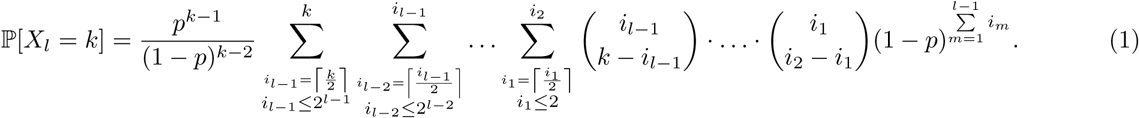

We hypothesise that this formula cannot be much simplified due to the multiplication of binomial coefficients. In Figure 1, we have run Monte Carlo simulations to generate draws from the PCR distribution. Comparing this simulation data to the corresponding pdf given by Eq. 1 yields very close agreement. However due to the nested sums, the pdf is very cumbersome to work with. We will therefore rather utilize the moments the PCR distribution generates.

**Fig. 1.**
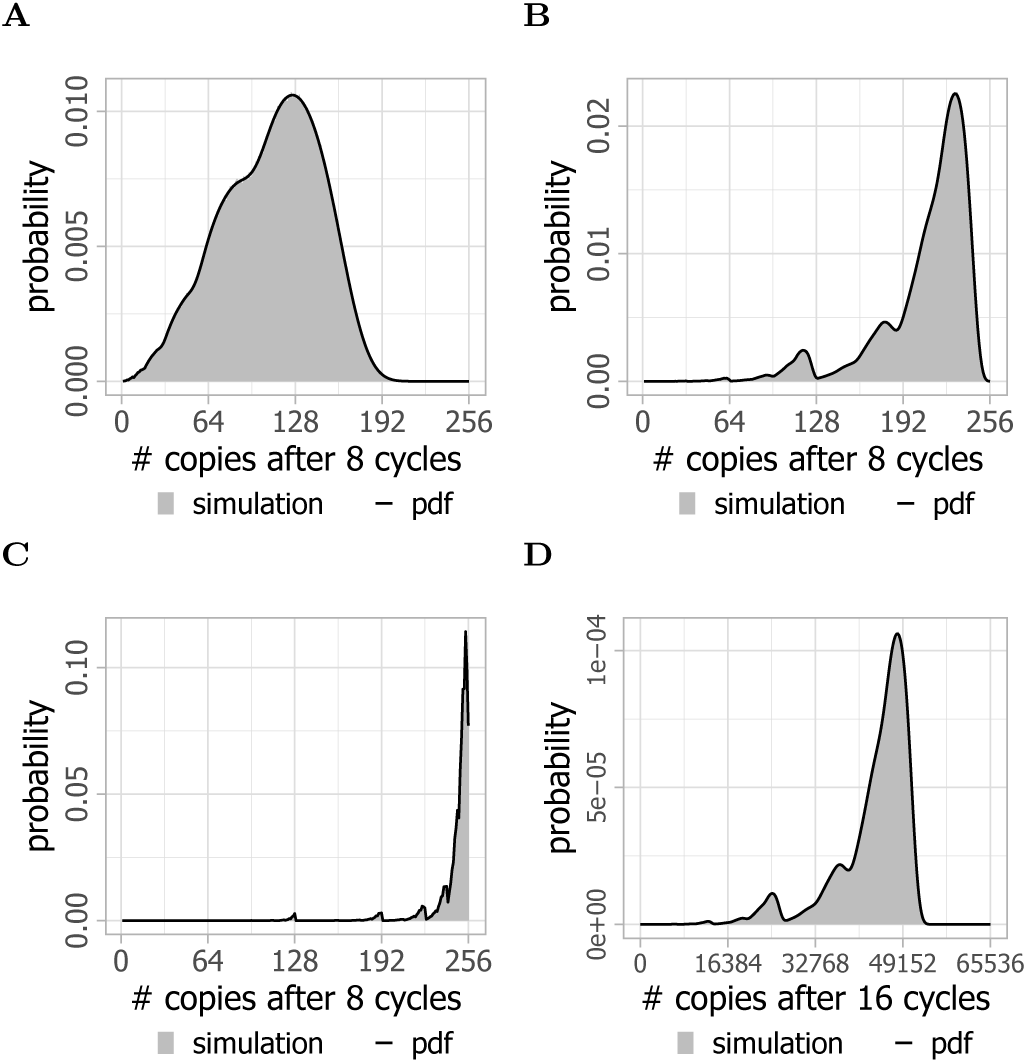
The pdf of the PCR distribution matches Monte Carlo simulations. For each plot, 10^6^ Monte Carlo simulations (blue) were performed to calculate the number of copies produced after **A**-**C** *l* = 8, **D** *l* = 16 cycles at PCR efficiency **A** *p* = 0.8, **B** *p* = 0.95, **C** *p* = 0.99, **D** *p* = 0.95. Additionally, the corresponding analytical pdf from Eq. 1 is plotted in black.

Despite the rather complicated form of the pdf in Eq. 1, the expectation and variance can be written in much simpler terms:

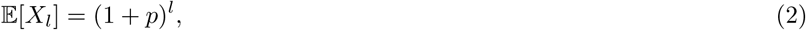

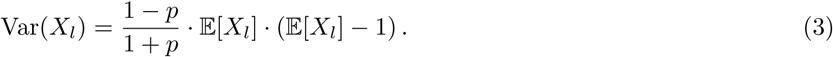

Interestingly, the formula for the expectation has been used for many decades to approximate the PCR efficiency in amplification experiments [17, 19, 28].

With the expectation and variance in hand, we can furthermore derive an equation for the coefficient of variation (CV) given by the ratio of the standard deviation to the expectation. The CV is an excellent measure for variability as it is comparable between random variables.

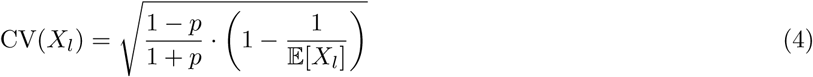

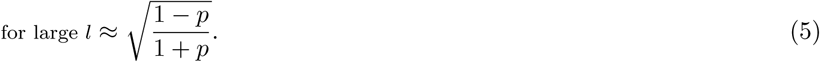

The CV increases with the number *l* of PCR cycles due to the term 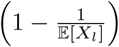. We note that this factor is between 0 and 1 and converges to 1 with increasing *l*. Therefore, the CV converges to the upper bound 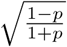. This upper bound is CV=0.229 for *p* = 0.9 and CV=0.071 for *p*=0.99. The speed of convergence depends on the PCR efficiency *p*. Performing more PCR cycles does not spoil the data by rising variabilty, but neither improves precision.

By setting a specific convergence threshold (see Methods), we interestingly find that the difference between nearly perfect PCR efficiency (*p* = 0.99) and a PCR efficiency of 0.8 is only one cycle, with 7 or 8 cycles required respectively to reach convergence (Figure S2). This implies that experiments with more than 8 cycles of PCR amplification can generally be considered to have a fixed CV that only depends on the PCR efficiency.

### Sequencing Probabilities

Let *K* denote the number of original mRNAs in the cell population, meaning on average

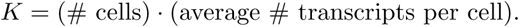

Each transcript *k* ∈ {1, …, *K*} produces *i*_*k*_ copies during PCR where each *i*_*k*_ is a realization drawn from the PCR distribution *X*_*l*_. After PCR amplification, all the PCR copies of all transcripts are sequenced together. Let *R* be the number of total PCR copies that is actually sequenced (i.e. the number of reads). Then each individual PCR copy has a chance 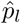 to be sequenced where 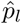 can be approximated by

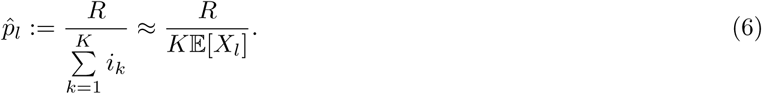

Eq. 6 can be perceived as an estimate for 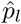 on the basis of the outcome of a specific experiment. For one particular original transcript *k*, the number of PCR copies actually sequenced out of the *i*_*k*_ copies produced is denoted by the random variable 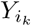. It follows a Poisson distribution as we are continuously drawing items from a large pool (on average 𝔼 [*X*_*l*_] copies for each of the *k* molecules) with an extremely low probability. Such a binomial distribution with large repetition and small probability per trial converges to the Poisson distribution. The corresponding Poisson parameter *λ*_*k*_ is given by 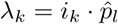.

In the setup described, we will observe an individual original molecule if at least one of its PCR copies is sequenced, meaning 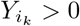. We call the probability to observe an original molecule *p*_*s*_. Then, we can show (see Methods) that

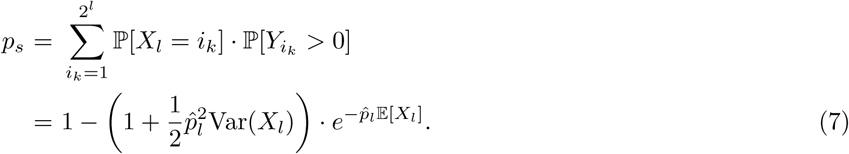

Utilizing our formulas for the moments of the PCR distribution (Eqs. 2, 3), we can express *p*_*s*_ in terms of the parameters of the experimental setup being the PCR efficiency *p*, the number of PCR cycles *l* and the probability to sequence a PCR copy 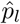:

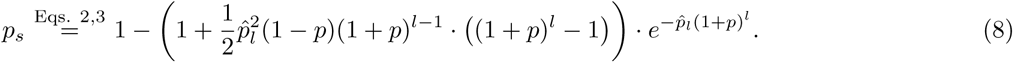

With the estimate from Eq. 6 for 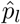, we can additionally display *p*_*s*_ to be:

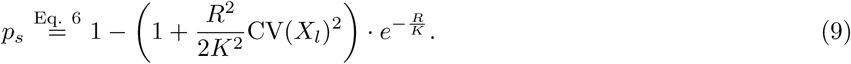

By Taylor expanding the exponential function to second order, we obtain

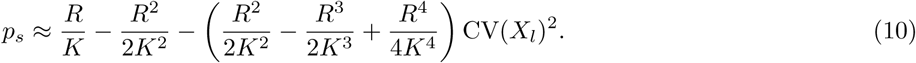

The variable *R* represents the total number of observed reads during sequencing (# reads). *K* is the total number of original transcripts in the cell population which is roughly given by (# cells) · (# transcripts per cell). We also stated that 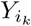 signifies the number of PCR copies actually sequenced for a specific transcript *k* ∈ {1, …, *K*}. Then, we can conclude

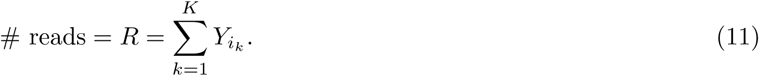

The number of reads *R* is then correlated to the number of PCR cycles *l* through the variables 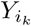

In order to obtain a simpler approximation for *p*_*s*_, we neglect the term 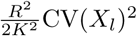 in Eq. 9. This simplifies the formula for *p*_*s*_ and we conclude

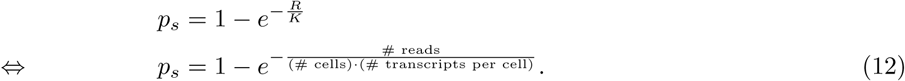

Eq. 12 provides a clear guideline on how to estimate *p*_*s*_ from experimental data. With these considerations, we can also call *p*_*s*_ the sequencing depth of the data set.

### Distribution of Observed Counts

So far, we have only considered what happens to one particular transcript of one specific gene that is originally present in one cell. We now investigate the distribution of the total number of transcripts for one particular gene *G*_1_ across multiple cells. Let *X*_in_ describe the input distribution of the number of transcripts of *G*_1_ found across a cell population and *X*_out_ be the corresponding output distribution, which represents the observed counts of *G*_1_ across the cell population after the sequencing experiment.

We now aim to relate the pdf of *X*_out_ to the pdf of *X*_in_. In the previous section, we have derived the probability *p*_*s*_ to sequence one particular transcript. We can utilize this result by considering the following: If in one particular cell there are *i* initial transcripts present of gene *G*_1_ and we observe *k* of them after the scRNA-seq experiment, it implies that we sequenced *k* transcripts and failed to sequence *i*−*k* transcripts. The number of possibilities to choose *k* out of these *i* transcripts is given by the binomial coefficient 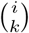. This leads to a binomial distribution with success probability *p*_*s*_ if the number of initial transcripts was known. Since this number is in fact not known, we need to sum over all probabilities for having *i* initial transcripts.

These considerations lead us to

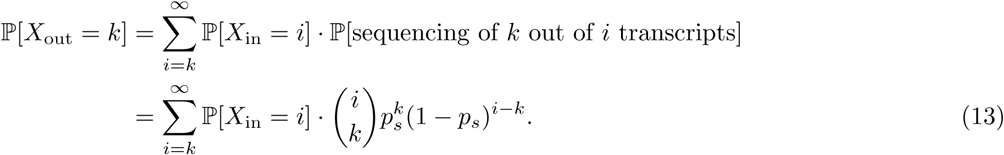

We can replace *p*_*s*_ by Eq. 8 to express the output distribution depending only on the input distribution, the PCR efficiency *p*, the number of PCR cycles *l* and the probability to sequence an individual PCR copy 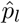, which are the parameters of the experimental setup. Given a specific input distribution, Eq. 13 completely characterizes the output distribution. It is typically assumed that *X*_in_ follows a log-normal distribution, *X*_in_ ∼ Log-normal(*µ, σ*^2^) [3], which will therefore be our choice for simulations.

It is possible to derive formulas for the expectation, variance and CV of *X*_out_ (see Methods Eqs. 41, 44, 45). However, in an experimental setting, we are rather interested in concluding characteristics of the input distribution when observing a certain output distribution. For these, we obtain:

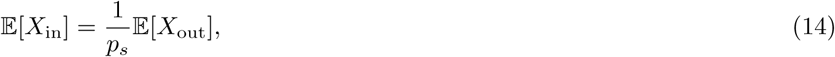

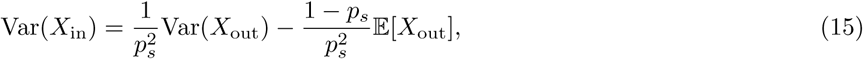

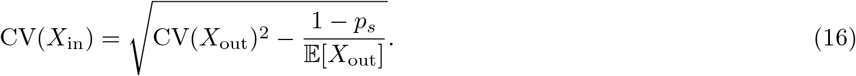

These moment relations apply with all input distributions. If we assumed a certain distribution type such as the log-normal distribution for the input distribution, then it would be completely defined by calculating these moments.

We can show that the CV of the output data is always larger than the CV of the input data in feasible cases:

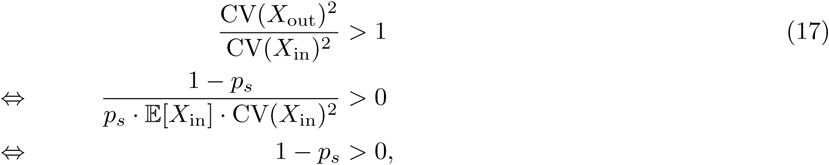

which is almost always true by definition. Only the case *p*_*s*_ = 1 does not satisfy this inequality. This would mean *X*_out_ = *X*_in_ which is experimentally not feasible. For *p*_*s*_ = 0, the inequality is not defined as it would imply *X*_out_ ≡ 0 so that no CV can be defined.

We can derive even more general statements from Eq. 13. It determines all moments of the output distribution for given moments of the input distribution (see Methods):

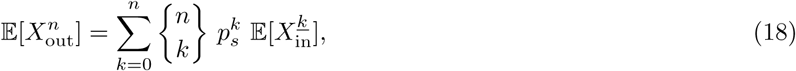

where 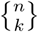 is the Sterling factor of second kind and 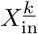 is the *k*-th falling power of *X*_in_ (see Methods Eq. 49). Vice versa, Eq. 18 can be rearranged to yield all moments of the input distribution from the output moments. We show this by an iterative approach which calculates the *n*-th moment of *X*_in_ from moments of *X*_in_ of order *n* − 1 and smaller and moments of *X*_out_ with highest order *n*:

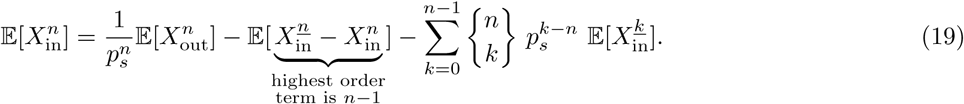

Knowledge of all moments implies a complete characterization of a distribution [16]. Hence, in principle we can recover the input distribution from scRNA-seq data, if the quality of the output data allows for calculation of higher moments.

### Mean-variance relationship in input and output distributions

While traditionally count data is modelled using Poisson distributions, sequencing count data is typically modelled using negative binomial distributions [1, 12, 27]. For scRNA-seq data, models relying on zero-inflated negative binomial distributions have become popular [9, 21, 25], although the presence of zero-inflation is still debated and doubtful [7, 29, 33]. Therefore, we will concentrate on the standard negative binomial distribution.

A Poisson distribution has the characteristic that its expectation *µ* and its variance *σ*^2^ are equal. For a negative binomial distribution with parameters *p* and *r*, the following relation between mean and variance holds:

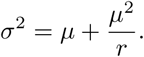

We can investigate both of these relations empirically in scRNA-seq data of HeLa cells taken from [30].

In Figure 3A, every point represents an individual gene. Clearly visible is the so-called overdispersion which makes the scRNA-seq data fit rather a negative binomial than a Poisson distribution. The reason for this overdispersion is often attributed to cell-to-cell variability (biological noise).

From our pdf for the output distribution in Eq. 13, we can already conclude that this is an accurate characterization. The presence of cell-to-cell variability is expressed by the fact that *X*_in_ is a distribution rather than a constant value. If we assumed the opposite, that *X*_in_ ≡ *n* ≥ *k*, then:

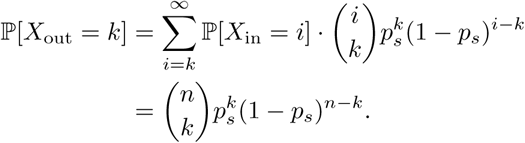

This is a binomial distribution with small success probability *p*_*s*_. We know that such a binomial can be approximated by a Poisson distribution. Hence in the absence of cell-to-cell variability, the sequencing count data would indeed follow a Poisson distribution.

For numerical simulations in Figure 2, we choose a log-normal distribution as our input distribution. For multiple parameter choices of the log-normal distribution, we generate observed counts as described in Figure S4. We also set *p*_*s*_ = 0.116 as we have estimated this for a particular scRNA-seq experiment involving HeLa cells (see Methods). We observe that the analytical output distribution matches the simulation data well (Figure 2). In addition to the analytical pdf, we also fit a Poisson and a negative binomial distribution. In all examples, it is possible to fit a negative binomial distribution very close to the analytical distribution, justifying its usage in numerical simulation. The Poisson fit on the other hand differs noticeably from the numerical simulation for high expression genes. The larger the CV of the input data, the larger the differences become. This is in line with our previous assertion that a constant input distribution would provide a Poisson output distribution. For lowly expressed genes on the other hand, the Poisson distribution provides a good fit to the data despite a large CV of the input data. The same fact can be derived from the mean-variance relation in Figure 3A.

**Fig. 2.**
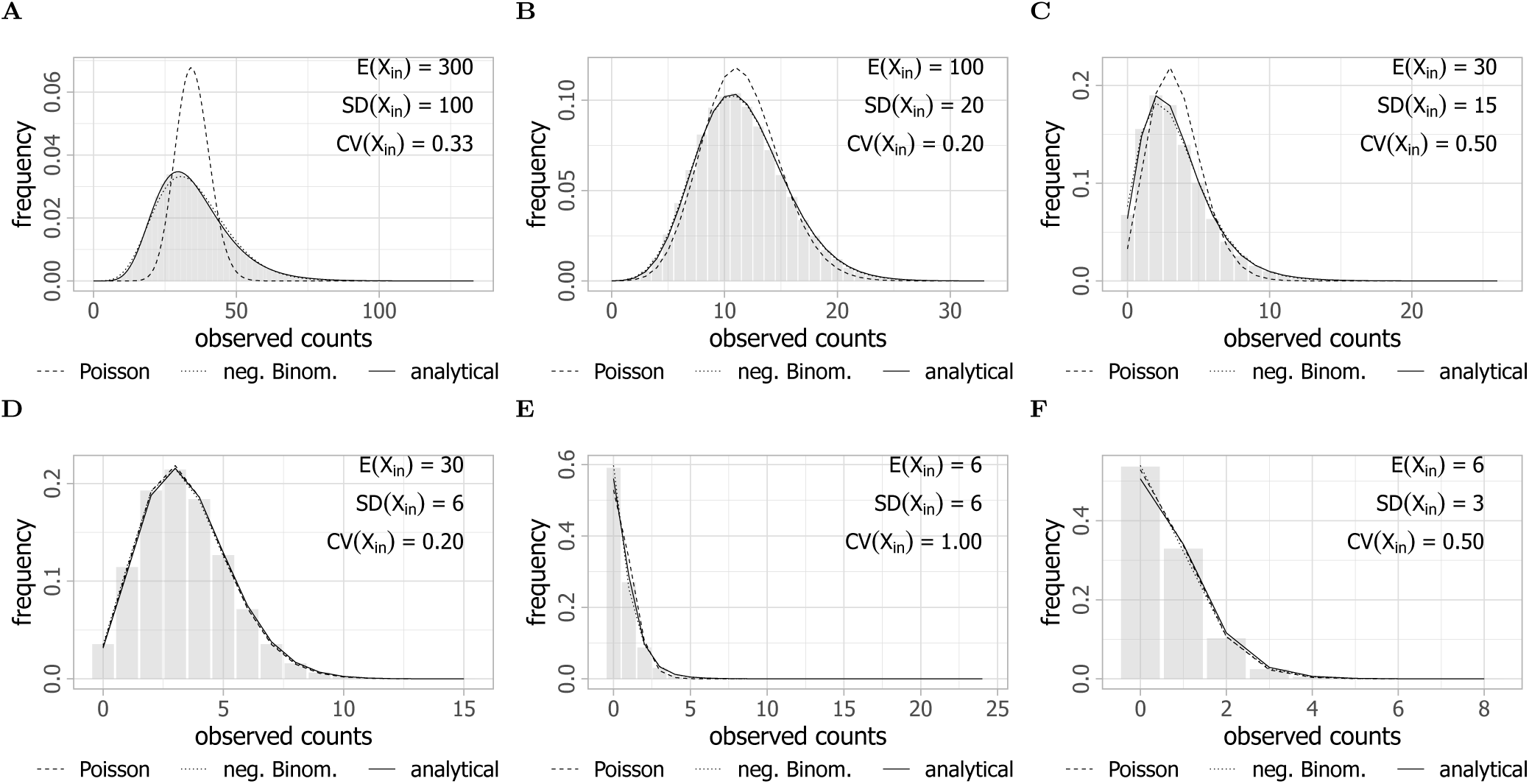
The analytical output distribution matches Monte Carlo simulations. In all cases, a negative binomial (blue) is fit to the data, as well as a Poisson distribution. Different input parameters are noted inside the plots.

**Fig. 3.**
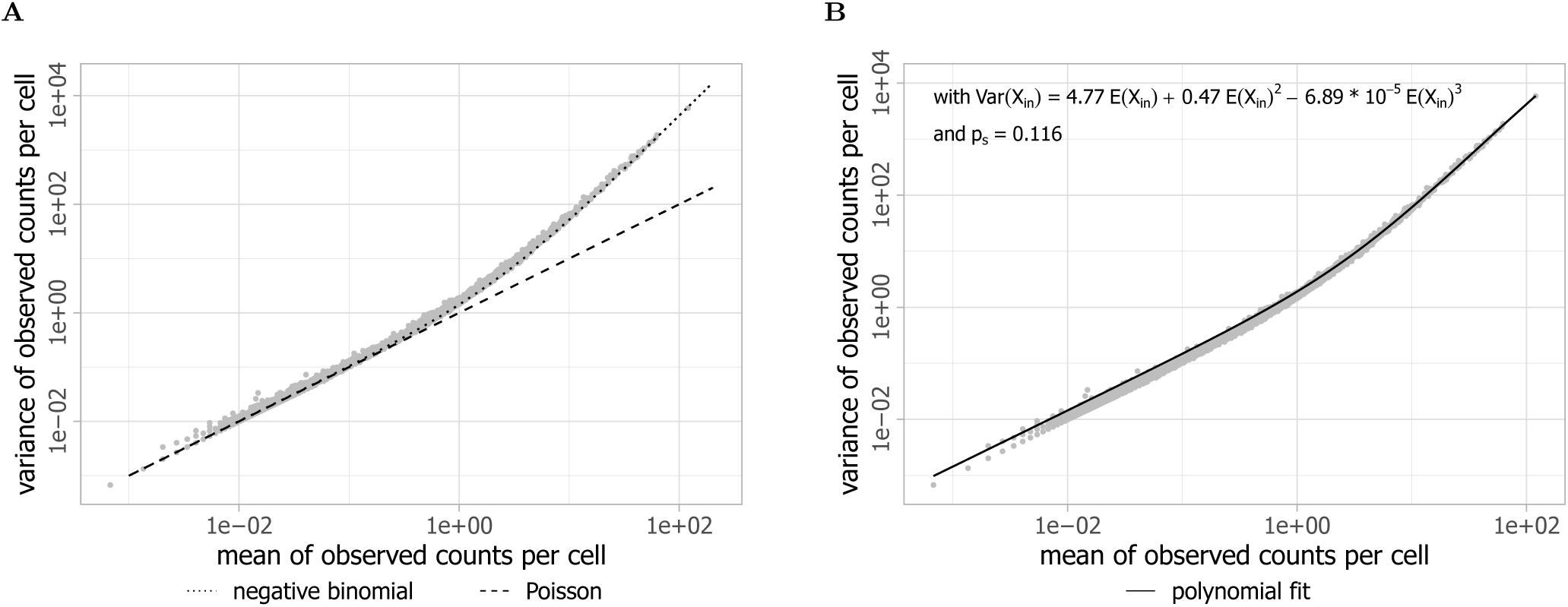
The mean-variance relation of scRNA-seq data for individual genes of a HeLa data set [30] satisfies a negative binomial rather than a Poisson distribution and establishes a relation between mean and variance of the input distribution. Each point represents an individual gene. Only non-variable genes with dispersion smaller than 0 are shown (see Methods of [30]). **A** A Poisson distribution (green) and negative binomial distribution (blue) are fitted to the data cloud. **B** By incorporating a cubic relation between the mean and variance of the input distribution, we find a good fit to the data. All three parameters of the cubic function are found to be significant for the fit.

The shape of the mean-variance relation in Figure 3A and other scRNA-seq data suggests that there is in fact a dependency of the variance on the expectation, i.e. a relation between moments exists. That entails the existence of a moment relation also for the input distribution. In Figure 3B, we assume a cubic dependency of the input variance on the input expectation

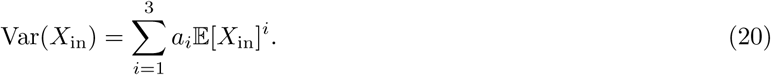

The resulting mean-variance relation fits the data reasonably well. If we furthermore assume the input distributions for the data to be log-normal [3], a relation between the two parameters *µ* and *σ*^2^ of a log-normal distribution can be established (see Methods). This means that if the output data shown in Figure 3 do stem from log-normal distributions, the parameters of these distributions have to satisfy the following equation:

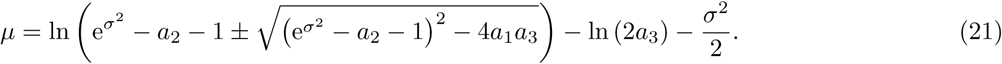

The parameters *a*_1_, *a*_2_ and *a*_3_ are fitted from the data and are displayed for the example data utilized here in Figure 3B. The quality of the fit suggests that we can specify the general assumption of a log-normal distribution for cellular transcripts to a log-normal distribution with a relation between the parameters like Eq. 21. The remaining parameter *σ* is fixed by the average transcript number.

One question of interest is to investigate how the shape of the input distribution compares to the output distribution. In order to compare these two distributions within one plot, we scale the input distribution by the factor *p*_*s*_ due to the relation of the expectations from Eq. 2: 𝔼 [*X*_out_] = *p*_*s*_ · 𝔼 [*X*_in_]. We observe in Figure 4 that the output distributions have fatter tails, while the input distributions are rather weighted towards its mean. This confirms the conclusion in Eq. 17 that the CV of the output distribution is larger than the input distribution.

**Fig. 4.**
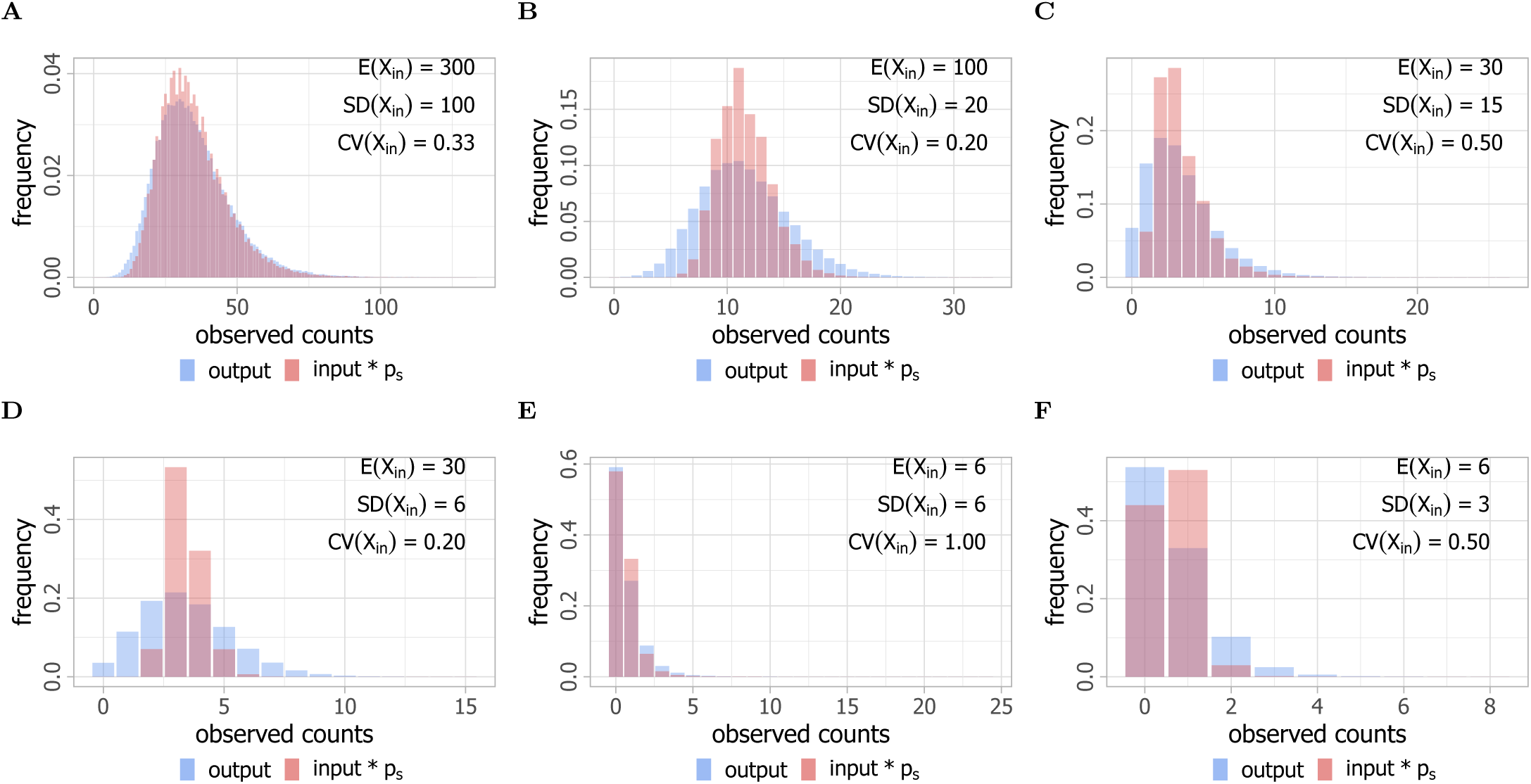
Visualization of the differences between input and output distribution. The histogram of the output distribution *X*_out_ (blue) of 10^5^ cells is plotted together with the histogram of the rescaled input values *p*_*s*_ *· X*_in_ (red) for *p*_*s*_ = 0.116 (see Methods). It illustrates that the CV of the output is always larger than the one of the input according to Eq. 17.

### Relation between input and output CV

From Eq. 16, we know that

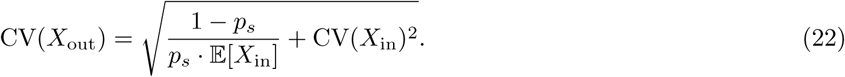

We investigate the dependence of CV(*X*_out_) on CV(*X*_in_) and the sequencing probability *p*_*s*_ (which we can also call the sequencing depth) in Figure 5. Most noticeably, when CV(*X*_in_) attains large values and thus dominates all other terms, the relation approaches a linear dependency. This is obvious also from Eq. 22. Considering the fold change of CV(*X*_out_) to CV(*X*_in_), we observe that small CV(*X*_in_) cause huge amplification of the variability whereas the fold change approaches 1 for larger values of CV(*X*_in_).

**Fig. 5.**
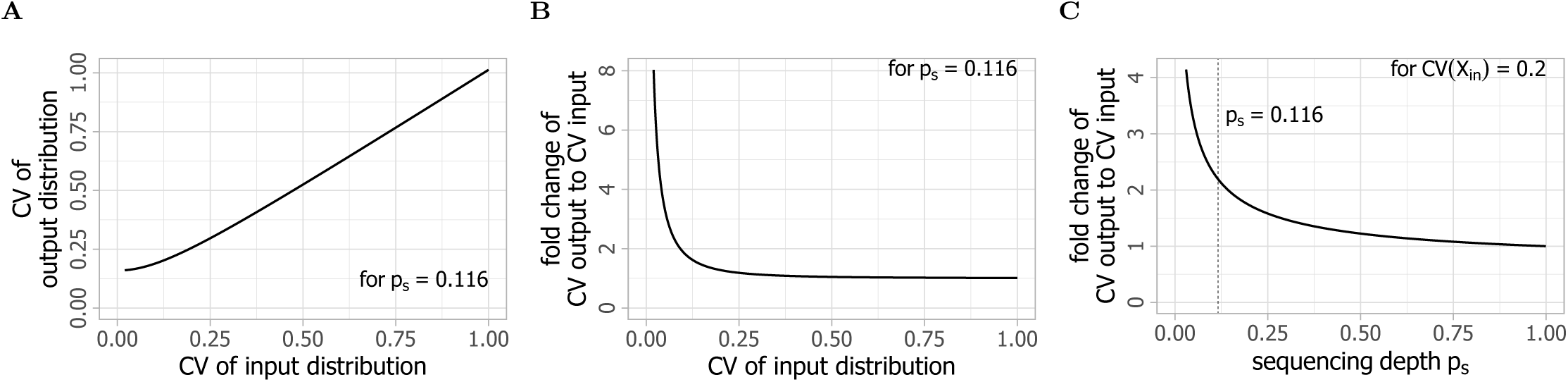
The CV of *X*_out_ shows characteristic dependencies on CV(*X*_in_) and *p*_*s*_ providing guidelines for obtaining desired noise levels. Effects of changes in input CV and success probability *p*_*s*_ on the output CV.

The effects of the sequencing depth *p*_*s*_ on CV(*X*_out_) are altogether not surprising as well. The more transcripts are sequenced, the smaller the increase of CV(*X*_out_). However, we do note that for the choice of CV(*X*_in_) = 0.2 and 𝔼 [*X*_in_] = 300 displayed here, the sequencing depth *p*_*s*_ = 0.116 is still located in an area of the hyperbola branch with increased growth. Therefore, increases in the sequencing depth *p*_*s*_ would still benefit noise reduction noticeably. Interestingly, these improvements start to drop off quickly (consider the slope of the graph). That means that improvements past a sequencing depth of around *p*_*s*_ = 0.5 would have far smaller effects on CV(*X*_out_). The same results are plotted in Figure S5 and Figure S6 in three dimensions for further illustration.

## Discussion

Simplified models do not capture every detail of an experiment and the precision of their predictions is limited. However given cell-to-cell variability, capturing all details limits the scope of models (and precision beyond the relative variance of noise is meaningless anyway). The advantage of simplified models is that they provide equations which can be used for a large class of experiments. In that spirit, we investigate the processes involved in scRNA-seq with a simplified model. It enables us to derive a set of equations useful for experiment design and analysis of results, and sheds light on some of the causes for and implications of various noise terms arising during scRNA-seq experiments.

Within the validity of our model setup, Eqs. 1, 2, 3, 8, 13 provide a complete characterization of a scRNA-seq experiment in terms of the experimental parameters PCR efficiency *p*, PCR cycle number *l* and the probability to sequence a PCR copy 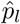. An experiment can be designed on that basis to provide a desired precision. The output distribution of an individual gene obeys Eq. 13 for a given input distribution. Knowledge of the moments of the input distribution independent of the distribution type suffice if we want to determine only moments of the output distribution. That is especially interesting in light of the finding that the input distribution obeys moment relations fixing its variance to the mean (Fig. 3, Eq. 20). Hence, both mean and variance of the output are determined by the mean of the input (see Eqs. 41, 44).

The purpose of scRNA-seq experiments usually is to determine the input distribution from the experiment’s output. We solve this inverse problem by calculating all moments of the input distribution from the moments of the output distribution by Eq. 19. Since in principle a distribution is completely determined if all its moments are known [16], this recovers the complete information of the input from the output distribution.

In practice, the output distributions are often not of sufficient quality allowing for the calculation of all moments. However, Eq. 19 enables us to determine as many input moments as we know output moments and thus guarantees that we can use all the information hidden in a measured output distribution. It is generally assumed that cellular RNA copy numbers obey a log-normal distribution [3]. That assumption can be verified on the basis of Eq. 19 if output distributions providing three or more moments are available.

We apply our results to a HeLa data set (Fig. 3). Two of the fundamental characteristics of single-cell data are that they exhibit relations between mean and variance and show higher variability than is typically expected for count data. The latter is called overdispersion (in comparison to a Poisson distribution). We determine the moment relation of the HeLa data set on the basis of Eqs. 41, 44 and obtain very good agreement with a third order polynomial (Fig. 3B). Assuming that the type of input distribution is the same for most of the genes (as Fig. 3A suggests to be the case) and that type to be log-normal, we determined the relation between the parameters of the cellular distribution characterising the HeLa data set. That is the complete characterization of the scRNA-seq input data which is possible within the validity of the assumptions since for each average number of RNA copies we can calculate the copy number distribution. The moment relation reproduces of course also the overdispersion in the HeLa data set. Relating it to Eq. 13 confirms overdispersion in general to be due to cell-to-cell variability, as has been assumed before.

The square root of the coefficient *a*_2_ of the moment relation of the input distribution (Eq. 20) is the coefficient of variation for highly expressed genes. The fit in Figure 3B shows this CV to be 0.686 for the HeLa data set. That means among the cells with transcript numbers within the range mean±SD we find a variability up to factor 5.4 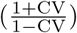 in transcript copy numbers, which we feel is surprisingly large for highly expressed genes.

We note that the assumptions we have made for the HeLa input distribution (log-normal distribution) is limited to homogeneous cell populations. Our formulas can in principal be applied to heterogeneous cell populations with bi- or multi-modal distributions just as well. The number of moments to be determined from the output distribution to recover the input distribution needs to be as large as the number of unknown parameters of the input distribution. If this is possible, appropriate ansatzes for the input distribution may also allow for calculating its parameters and thus to determine bi-modal input as well.

The results of our study can be used in several ways. They help to optimize the experimental parameters and they can be used to obtain as much information on the input distribution moments as desired and supported by the quality of the output data. It is a theory study which we hope will be picked up and checked by experimental labs.

## Acknowledgements

We would like to thank Laleh Haghverdi for very helpful comments and advice on the manuscript.

## Methods

### The probability distribution function caused by PCR

We calculate

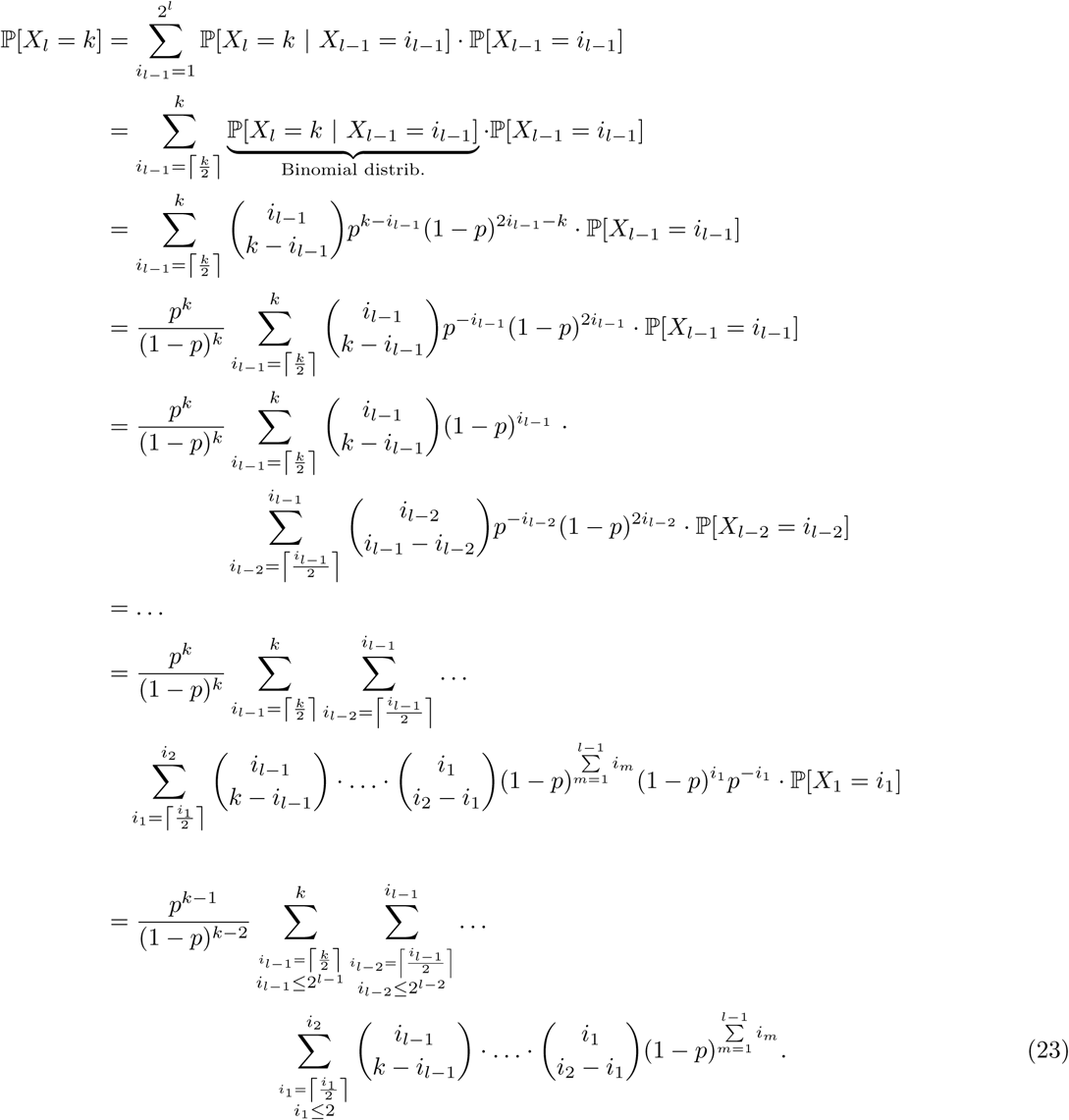

With some re-indexing, we can show that this is equivalent to

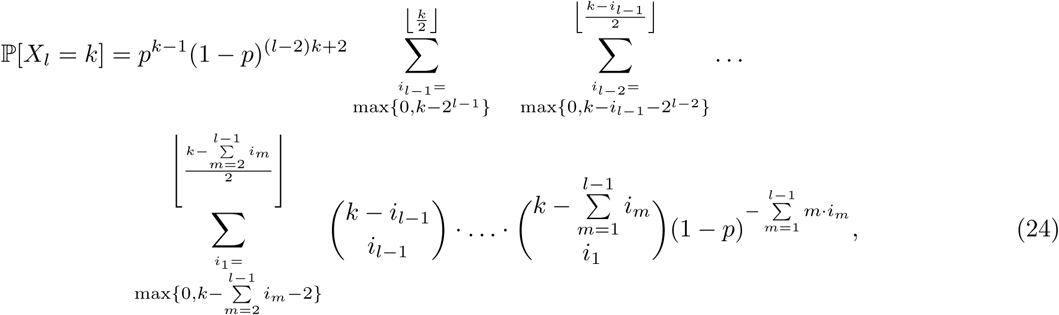

which is also equivalent to

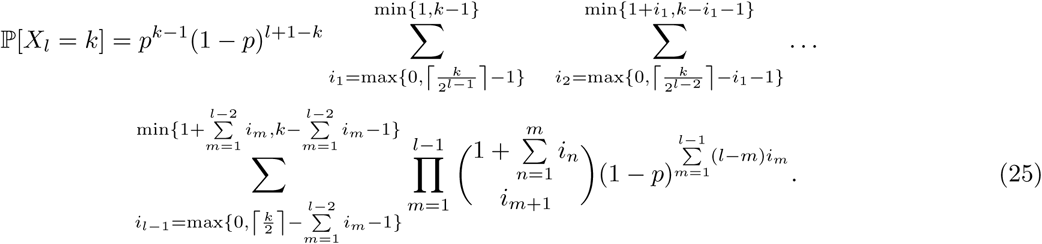

### Proof for the moments of the PCR distribution

We now prove the Eqs. 2, 3 with the help of the induction principle.

### Expectation

We stated previously that the increase in number of copies from *X*_*l*−1_ to *X*_*l*_ follows a binomial distribution (*X*_*l*_ − *X*_*l*−1_|*X*_*l*−1_ = *n*) ∼ Binom(*n, p*). We also note that for any binomial distribution *Z* ∼ Binom(*n, p*), it holds that 𝔼 [*Z*] = *n* · *p*. Hence,

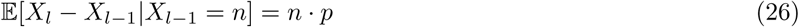

and

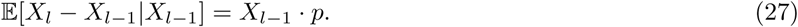

We now perform a proof via induction.

*Base case: l* = 1

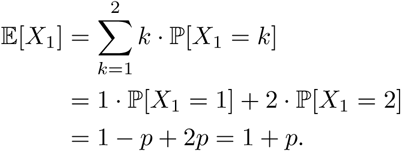

*Induction hypothesis:* 𝔼 [*X*_*l*−1_] = (1 + *p*)^*l*−1^.

*Induction step:*

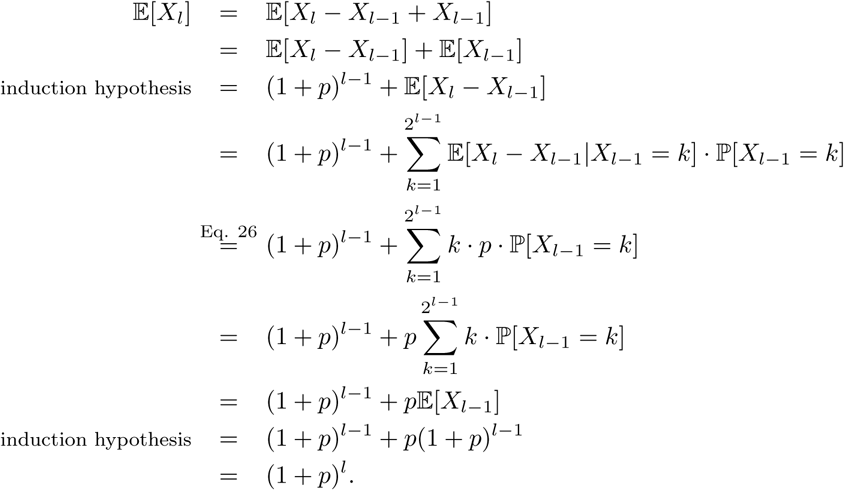

□

Hence, the expectation is calculated with the help of the expectation of the output distribution.

### Variance

From (*X*_*l*_ −*X*_*l*−1_|*X*_*l*−1_ = *n*) ∼ Binom(*n, p*), we can also conclude that Var(*X*_*l*_ − *X*_*l*−1_ | *X*_*l*−1_ = *n*) = *p* · (1 − *p*) · *n* and furthermore

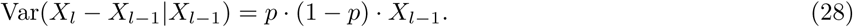

The law of total variance states for two random variables *X* and *Y* where *X* has finite variance that

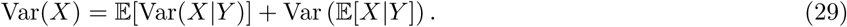

For the proof of the variance formula in Eq. 3, we need to assess the variance of the increase of copy numbers between two cycles Var(*X*_*l*_ − *X*_*l*−1_):

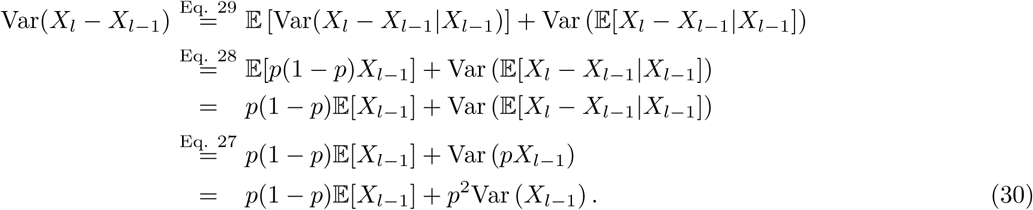

We also require an expression for the covariance of *X*_*l*_ − *X*_*l*−1_ and *X*_*l*−1_:

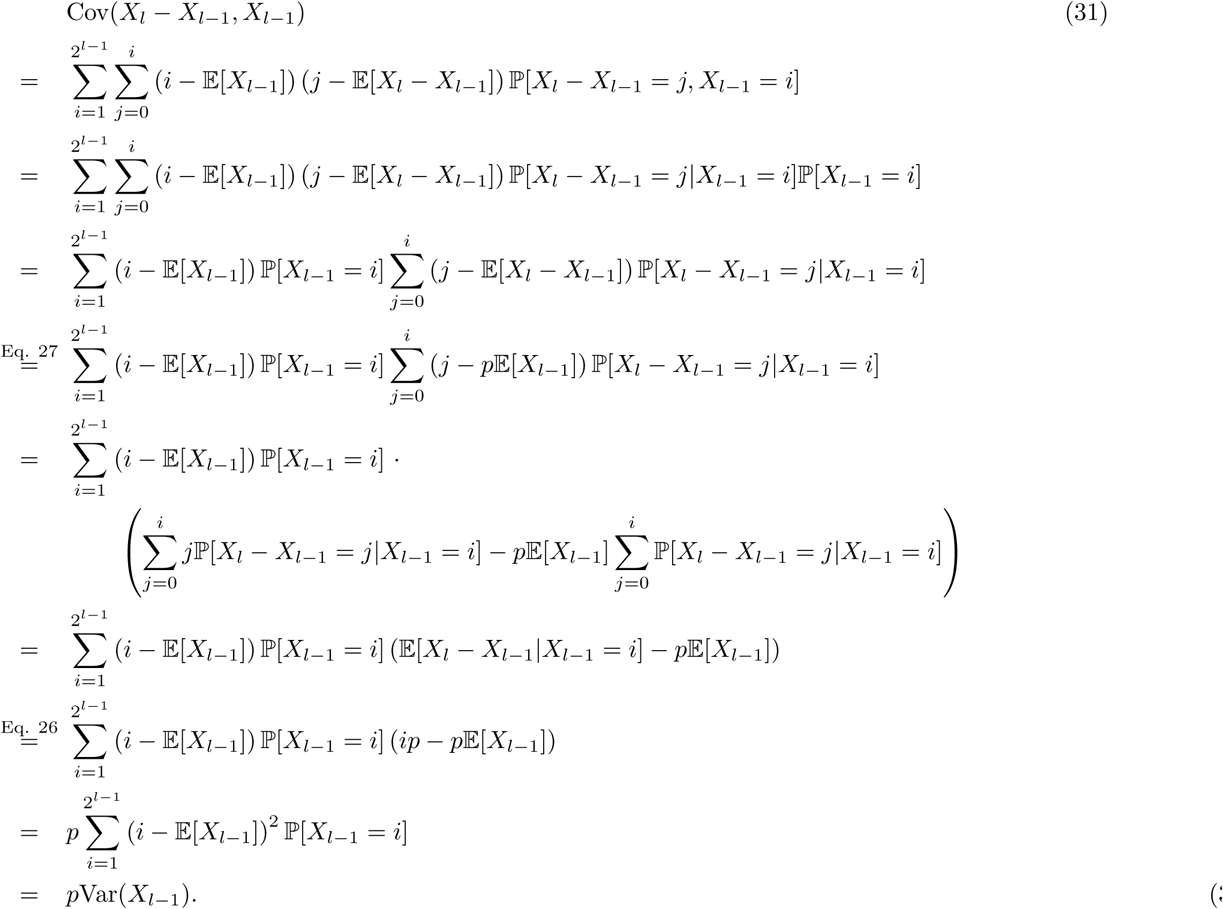

Now, we can proof Eq. 3 via induction:

*Base case: l* = 1

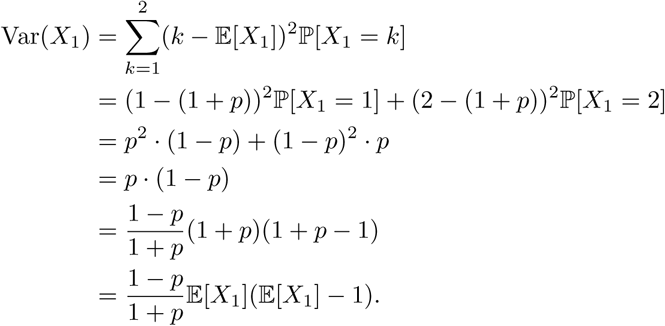

*Induction hypothesis:* 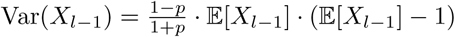.

*Induction step:*

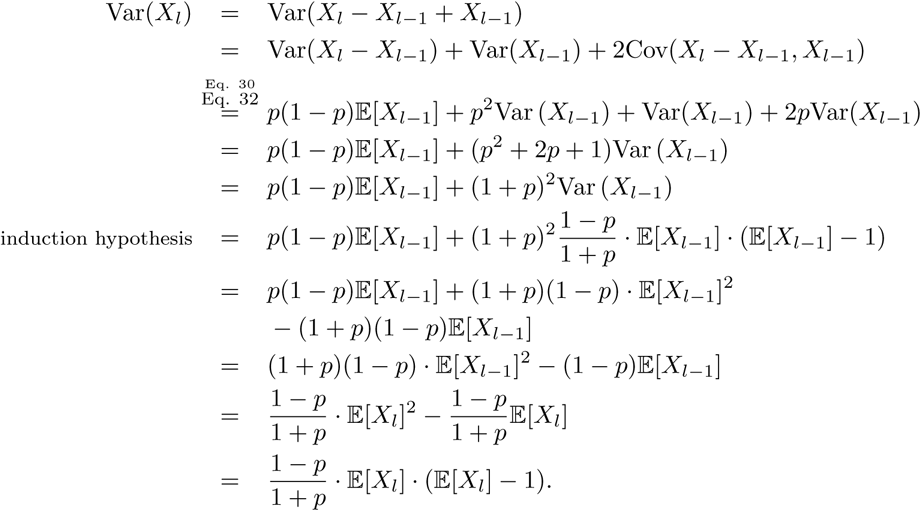

□

### Convergence of the coefficient of variation of the PCR distribution

We define that the term 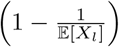 from Eq. 4 is negligible when 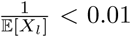 since then 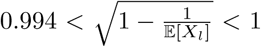 meaning the actual CV deviates less than 0.6% from the approximation. This essentially removes the dependency of the CV on the number of cycles *l*. We call the term 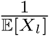 the CV convergence factor which should be large when *l* is small and approach 0 when *l* is large.

In Figure S2, we observe a sharp drop of the CV convergence factor between 1 and 7 cycles. We have coloured all values 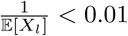 white so as to investigate how many PCR cycles are necessary to saturate the CV at different PCR efficiencies.

### Simulation data for the PCR distribution matches formulas for moments

In addition to the visual confirmation of our calculations in Figure 1, we can also investigate the mean *µ*, variance *σ*^2^ and CV of the simulation data and compare them to our analytical formulas in Eqs. 2, 3, 5 for *X*_*l*_:

| | $\mu$ | $\mathbb{E}[X_l]$ | $\sigma$ | $\text{SD}(X_l)$ | $\sigma^2$ | $\text{Var}(X_l)$ | CV | $\text{CV}(X_l)$ |
| --- | --- | --- | --- | --- | --- | --- | --- | --- |
| $l = 8,$<br>$p = 0.8$ | 110.18 | 110.20 | 36.55 | 36.57 | 1336 | 1337 | 0.332 | 0.332 |
| $l = 8,$<br>$p = 0.95$ | 209.12 | 209.06 | 33.37 | 33.40 | 1114 | 1115 | 0.160 | 0.160 |
| $l = 8,$<br>$p = 0.99$ | 245.92 | 249.94 | 17.43 | 17.40 | 304 | 303 | 0.071 | 0.071 |
| $l = 16,$<br>$p = 0.95$ | 43713 | 43707 | 6995 | 6999 | $48 \cdot 10^6$ | $48 \cdot 10^6$ | 0.160 | 0.160 |

The data are within a very reasonable error margin and therefore confirm our analytical conclusions.

### Markov chain formulation of the PCR distribution

The problem of PCR amplification can also be posed as a Markov chain illustrated for *l* = 2 in Figure S3. We note that for larger *l* the stationary probabilities for state 3 and 4 will have to be adjusted. These are set to 1 for *X*_2_ in order to adhere to the rule that the rows of transition probabilities have to sum to 1. We can state the corresponding probability transition matrix 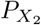 which is given by

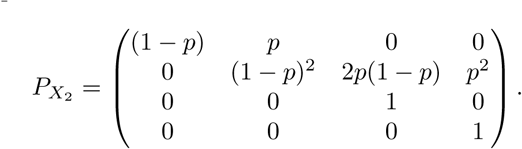

The pattern that emerges is that this is an upper triangle matrix. Within each row *i*, all entries (*i, j*) with *j* < *i* equals 0. The diagonal element (*i, i*) is the starting point of writing down all elements of a binomial distribution for success *p* and failure (1 − *p*) followed by zeros until the end of the row is reached. This is only true for the first 2^*l*−1^ rows. The bottom half of the matrix is the identity matrix. In more general terms, we have the following pattern:

- (*i, j*) = 0, if *j* < *i*,
- (*i, i*) = (1 − *p*)^*i*^, if *i* ≤ 2^*l*−1^,
- (*i, i*) = 1, if *i* > 2^*l*−1^,
- 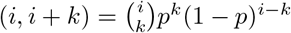, if 0 < *k* < 2*i* and *i* ≤ 2^*l*−1^,
- (*i*, 2*i*) = *p*^*i*^, if *i* ≤ 2^*l*−1^,
- (*i, j*) = 0, if *j* > 2*i*,
- (*i, j*) = 0, if *j* > *i* and *i* > 2^*l*−1^.

A general probability transition matrix for *X*_*l*_ takes the form:

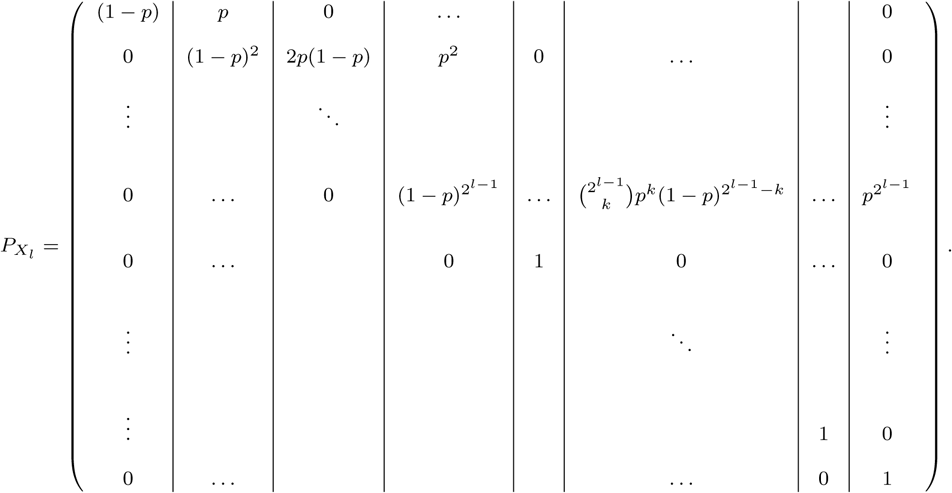

From the theory of Markov chains, we know that if we consider *l* steps along the Markov chain, the transition probabilities would be provided by the *l*-th power of 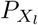. Furthermore, the index (*i, j*) of that power yields the probability to transition from state *i* to state *j* in *l* steps. Since we are only interested in the transition probabilities from starting state 1, we can conclude that

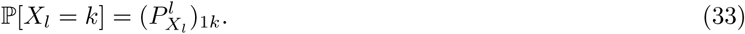

This means that the probability to obtain *k* copies after *l* PCR cycles is given by the entry (1, *k*) of the matrix obtained by raising 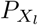 to the *l*-th power.

### Probability to observe a molecule

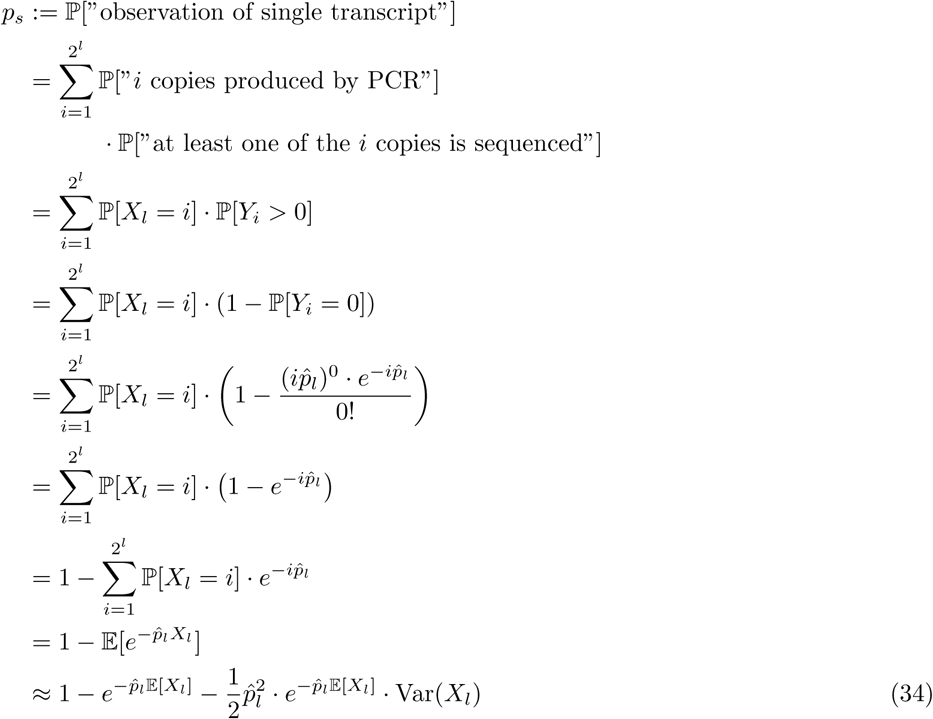

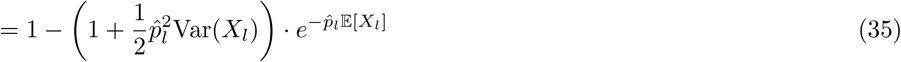

For Eq. 7 to hold, we need to prove the approximation in Eq. 34.

### Proof of approximation

For a general infinitely differentiable function *f* (*X*) of a random variable *X*, we can consider the Taylor expansion around the mean:

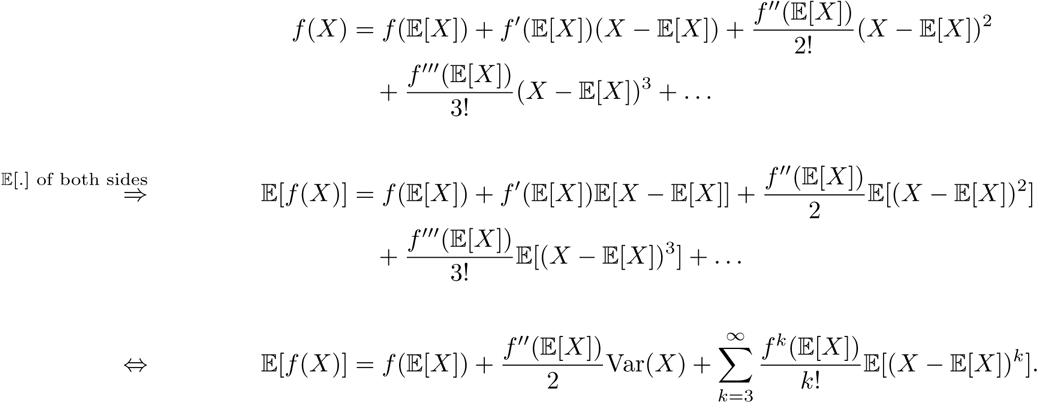

We now choose 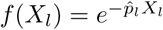 and note

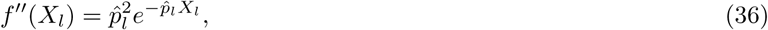

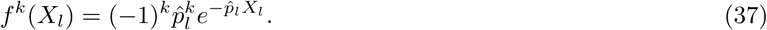

Hence,

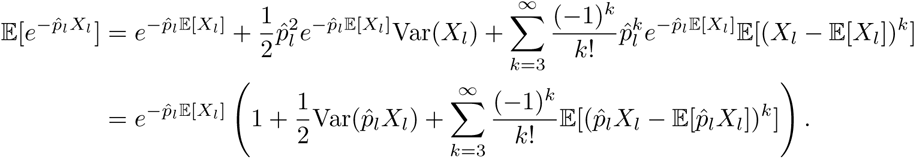

The elements of the sum are declining rapidly if 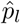 is small. Since the series is moreover alternating, it is reasonable to ignore terms for *k* ≥ 3.

### Estimating parameter values for 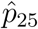 and *p*_*s*_

We want to roughly estimate 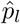 from experimental parameters. We have

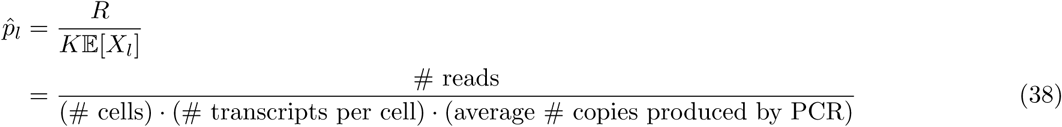

where # reads is the total number of observed reads during sequencing which we previously denoted by *R, K* is the average total number of original transcripts in the cell population and 𝔼 [*X*_*l*_] is the average number of PCR copies produced per transcript. We also stated that 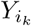 signifies the number of PCR copies actually sequenced for a specific transcript *k* ∈ {1, …, *K*}. Then, we can conclude

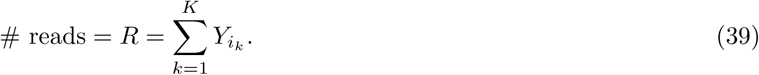

The number of reads depends on the number of PCR cycles *l*.

It is estimated that a single cell contains about 1 million mRNA transcripts. In an experiment involving HeLa data [30], we perform around *l* = 25 PCR cycles at an estimated *p* = 0.95 PCR efficiency. The sequencing run for the HeLa data yields roughly 2 · 10^8^ mapped reads for 1624 cells (before filtering). We therefore estimate the sequencing efficiency 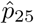 to be

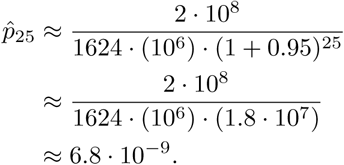

For this particular data set, we can then also estimate the probability *p*_*s*_ to detect a specific transcript with Eq. 12:

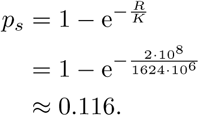

### Moments of Output Distribution

In order to characterize the output distribution in Eq. 13, we investigate the first and second moments as well as the CV.

### Expectation

The expectation is calculated as

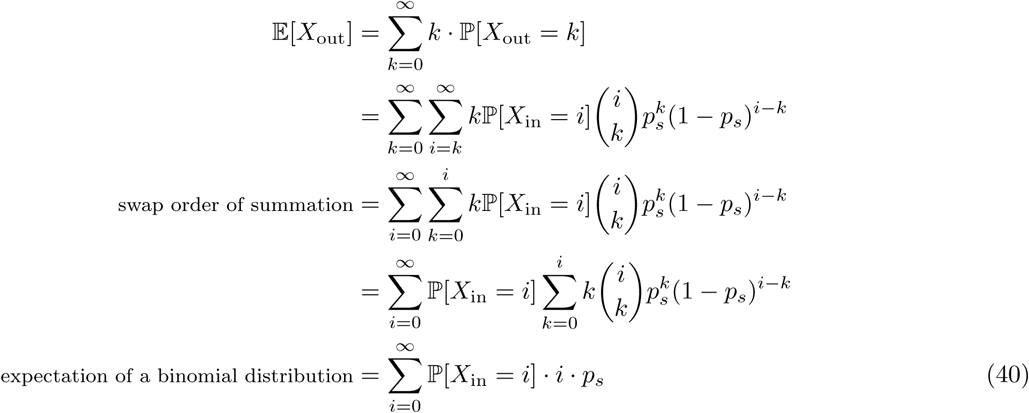

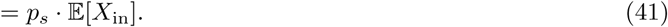

### Variance

We first consider the second moment

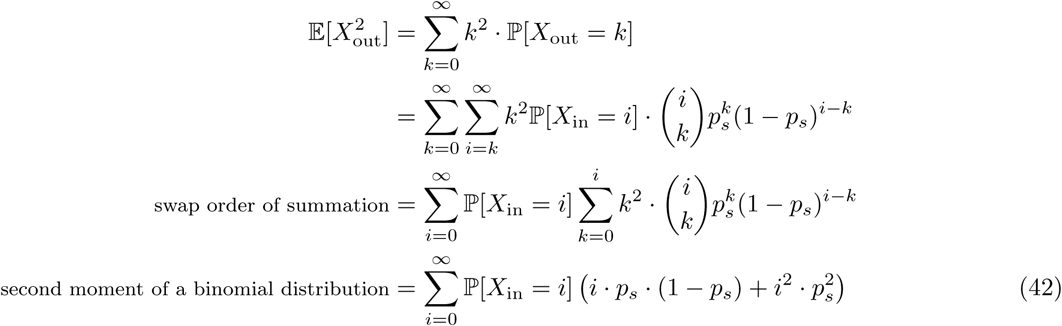

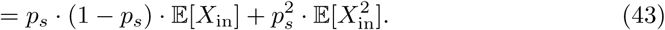

Now, the variance is given by

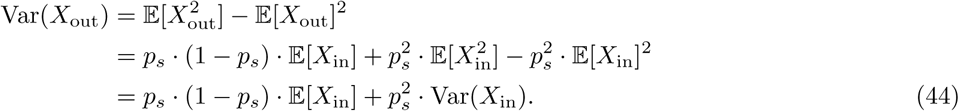

### Coefficient of Variation

The coefficient of variation is obtained by considering

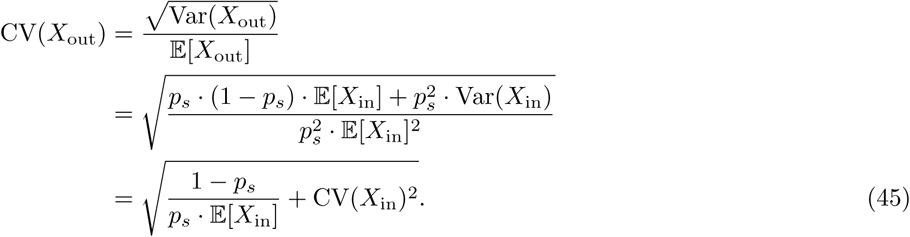

### General moments

We can generalize the previous considerations and write down an equation for the *n*-th moment of *X*_out_. Then,

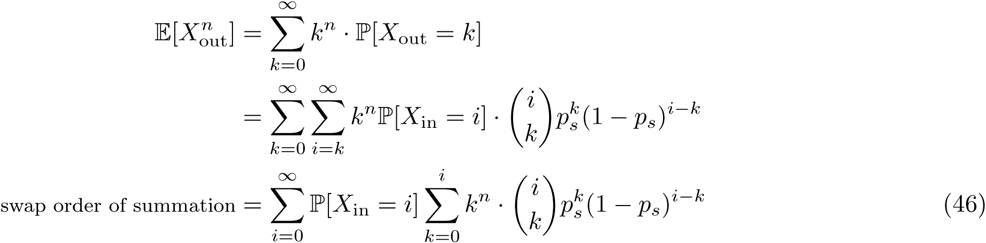

The second sum corresponds to the *n*-th moment of a binomal distribution Binom(*i, p*_*s*_) as given by [18]:

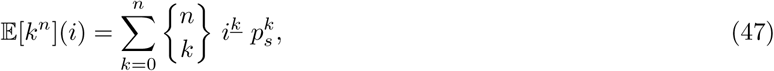

where 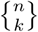 is the Stirling number of the second kind given by

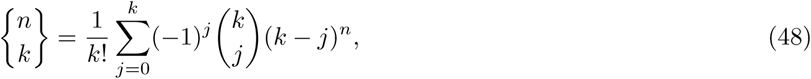

and *i*^*<u>k</u>*^ is the *k*-th falling power (also called falling factorial) of *i* which is calculated as

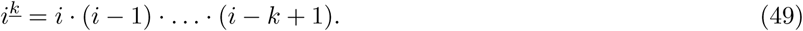

Then from Eq. 46, we conclude

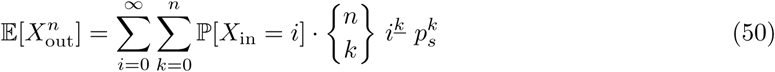

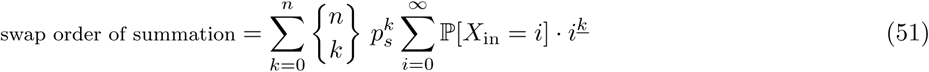

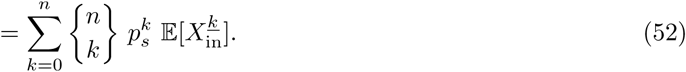

This provides a straightforward equation for calculating the *n*-th moment of the output distribution which depends only on the first *n* moments of the input distribution. We can check our previous calculations from Eqs. 41, 43:

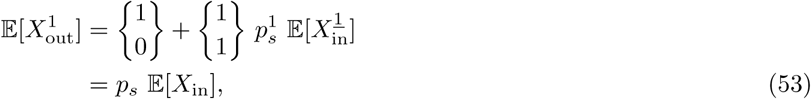

and

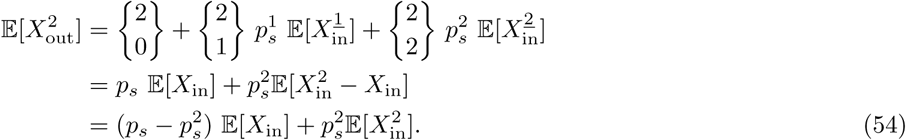

The third moment is obtained by considering

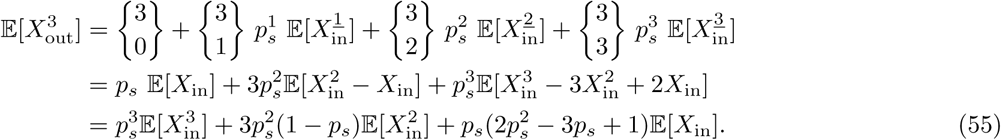

We note that the highest order moment of *X*_in_ required to calculate the *n*-th moment of *X*_out_ is also the *n*-th moment. This enables an iterative strategy for calculating the moments for *X*_in_ given the moments of *X*_out_. If all moments of *X*_out_ are known, the distribution of *X*_in_ including its distribution type is fully defined.

Eq. 52 can be rearranged to obtain an expression for 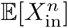 that only depends on the first *n* moments of *X*_out_ and the first *n* − 1 moments of *X*_in_:

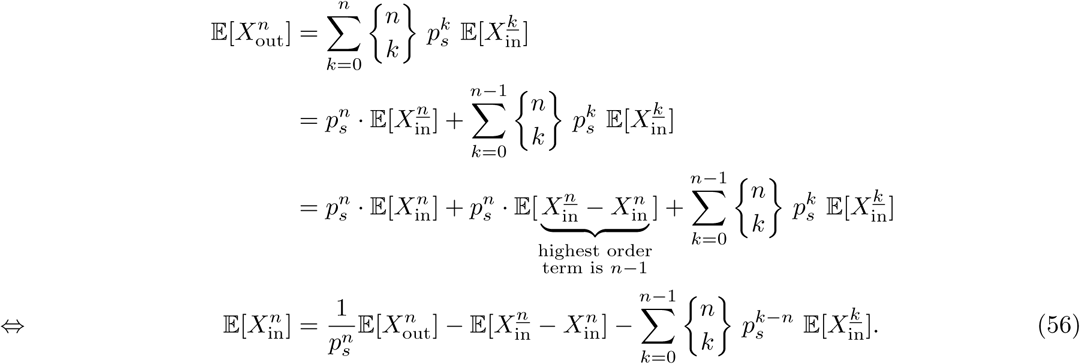

All terms on the right hand-side can be iteratively expressed by moments of the output distribution. If all moments of *X*_out_ exist, so do the moments of *X*_in_ which means that the probability distribution of *X*_out_ completely defines the probability distribution of *X*_in_.

### Relations of the parameters of a log-normal distribution required to satisfy the mean-variance relation of experimental scRNA-seq data

In Figure 3A, there is a clear dependency of the variance on the expectation of the output data of a scRNA-seq experiment. We have fitted them with a Poisson and a negative binomial distribution. We now utilize the output distribution we have derived and state requirements on the parameters of the log-normal distributions which is the distribution type typically chosen to model input distributions.

From Eqs. 41, 44, we have

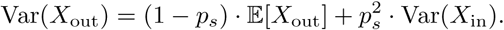

Let *X*_in_ satisfy the following relation:

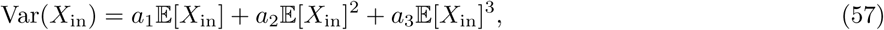

where *a*_1_, *a*_2_ and *a*_3_ are parameters. Hence, we assume a polynomial relation between the expectation of the input distribution and its variance. This yields

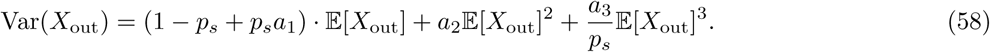

We can fit the parameters and obtain the graph shown in Figure 3B. We find all three parameters to be significant, meaning all three orders of the polynomial are essential for the fit.

If we assume *X*_in_ ∼ ℒ 𝒩(*µ, σ*^2^), then we can establish a relation between the two parameters *µ* and *σ*^2^ of the input distribution with the help of Eq. 57. For a log-normally distributed variable *Z*, we know

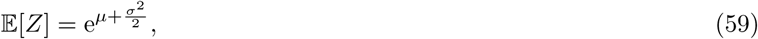

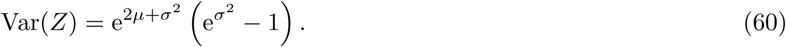

Then, we conclude from Eq. 57 that

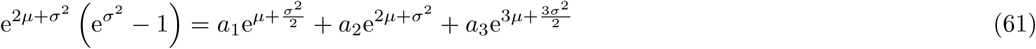

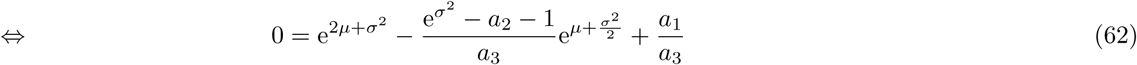

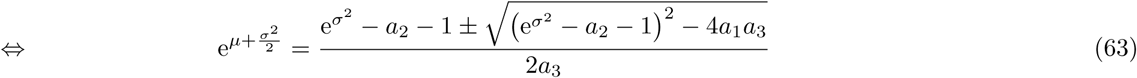

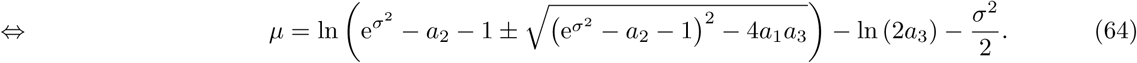

Thus, the log-normal distributions which we assume to generate the output distributions shown in Figure 3A and Figure 3B should satisfy Eq. 64 with the parameters *a*_1_, *a*_2_ and *a*_3_ fitted in Figure 3B.

## 2 Supplementary figures

**Fig. S1.**
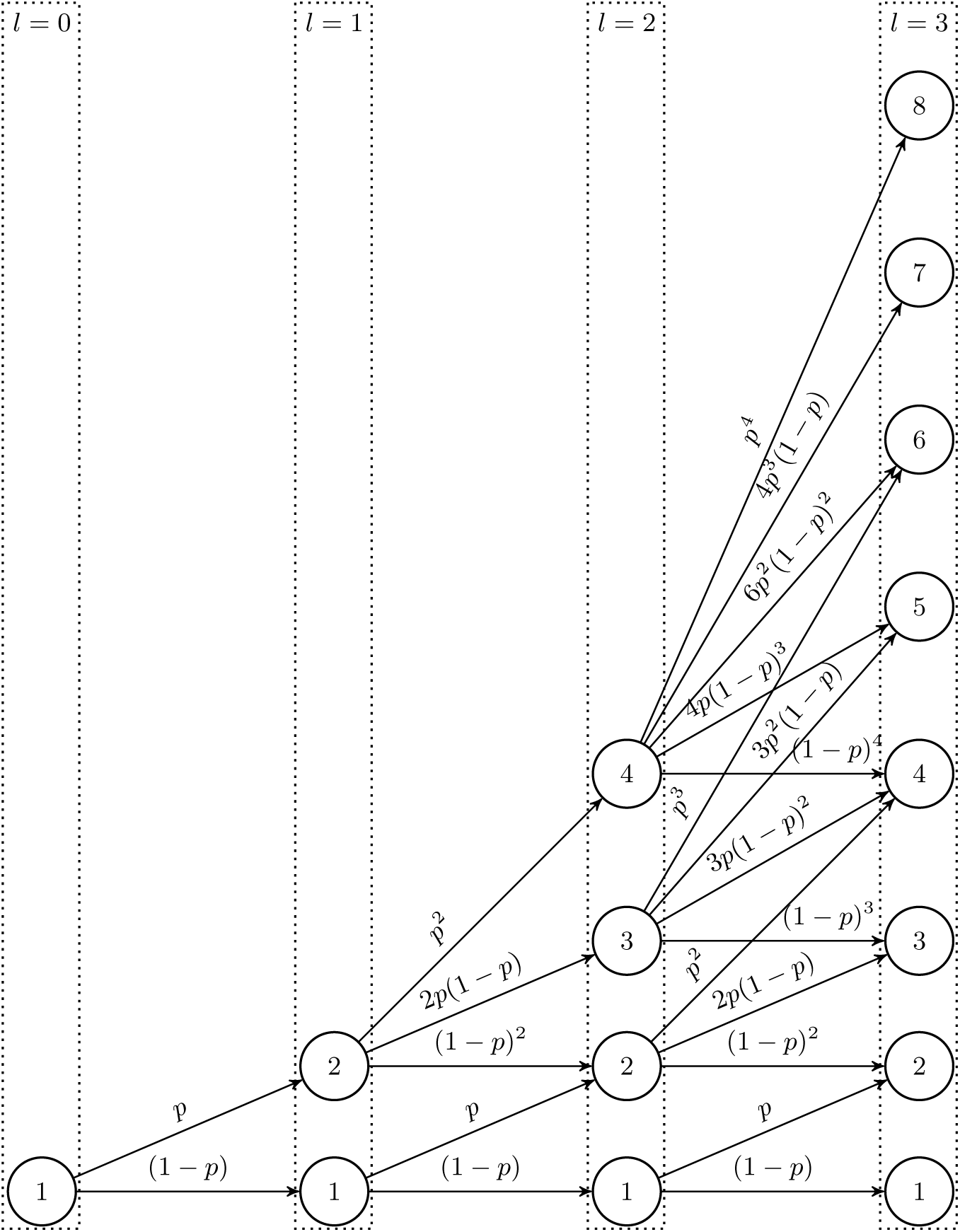
Probability transitions for the first three cycles of PCR. Each node represents the number of PCR copies produced after *l* cycles where *l* is represented by the columns.

**Fig. S2.**
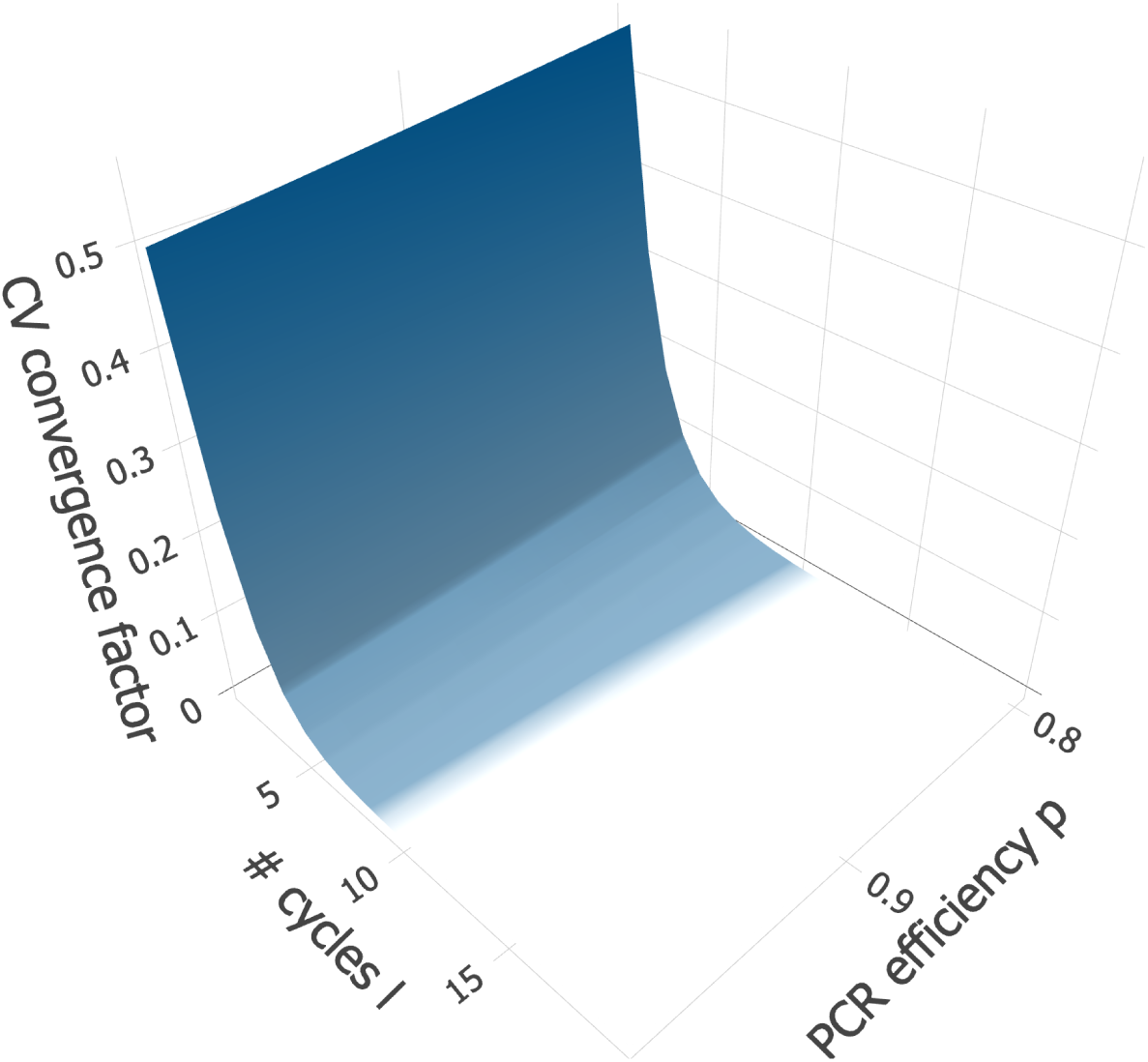
The CV convergence factor converges fast, reaching the convergence threshold at around 7 and 8 cycles. The CV convergence factor is calculated in dependence on parameters *l* and *p*.

**Fig. S3.**
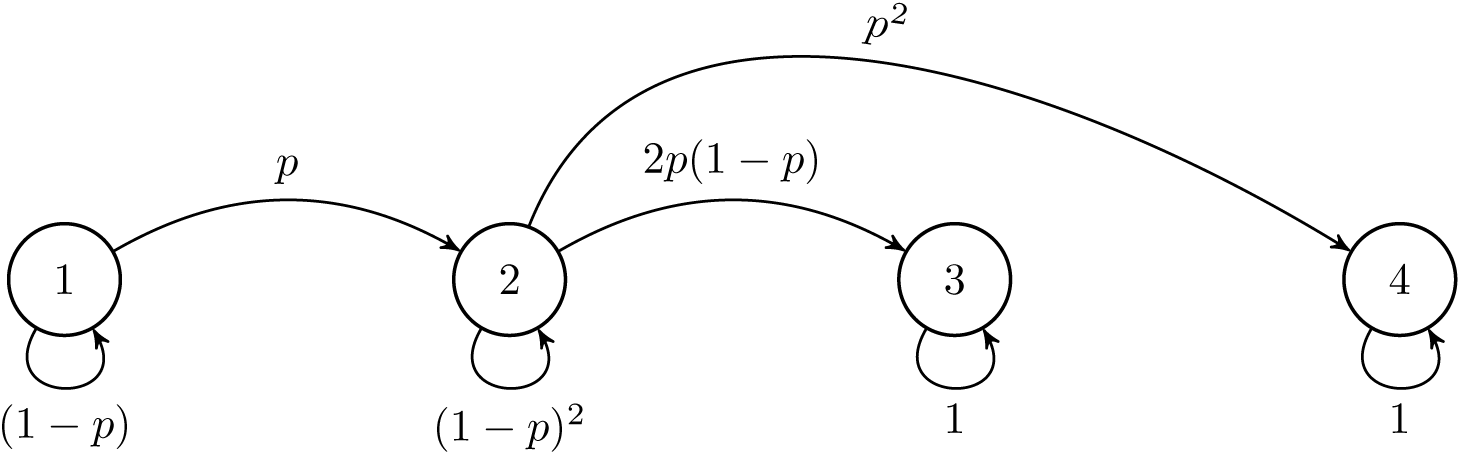
The PCR distribution as a Markov chain.

**Fig. S4.**
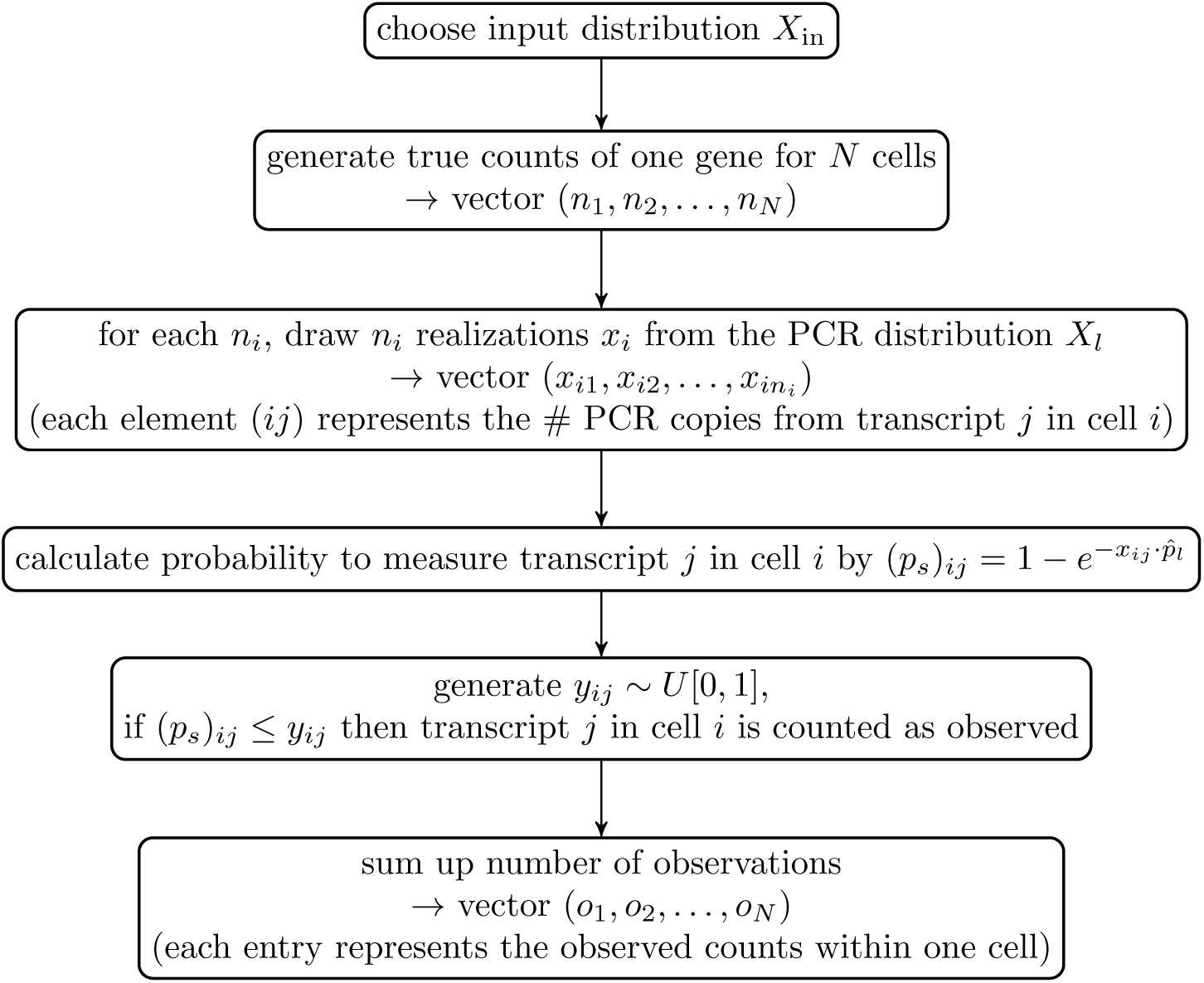
Principle steps of a numerical simulation of a simplified scRNA-seq experiment to obtain observed counts given an input distribution.

**Fig. S5.**
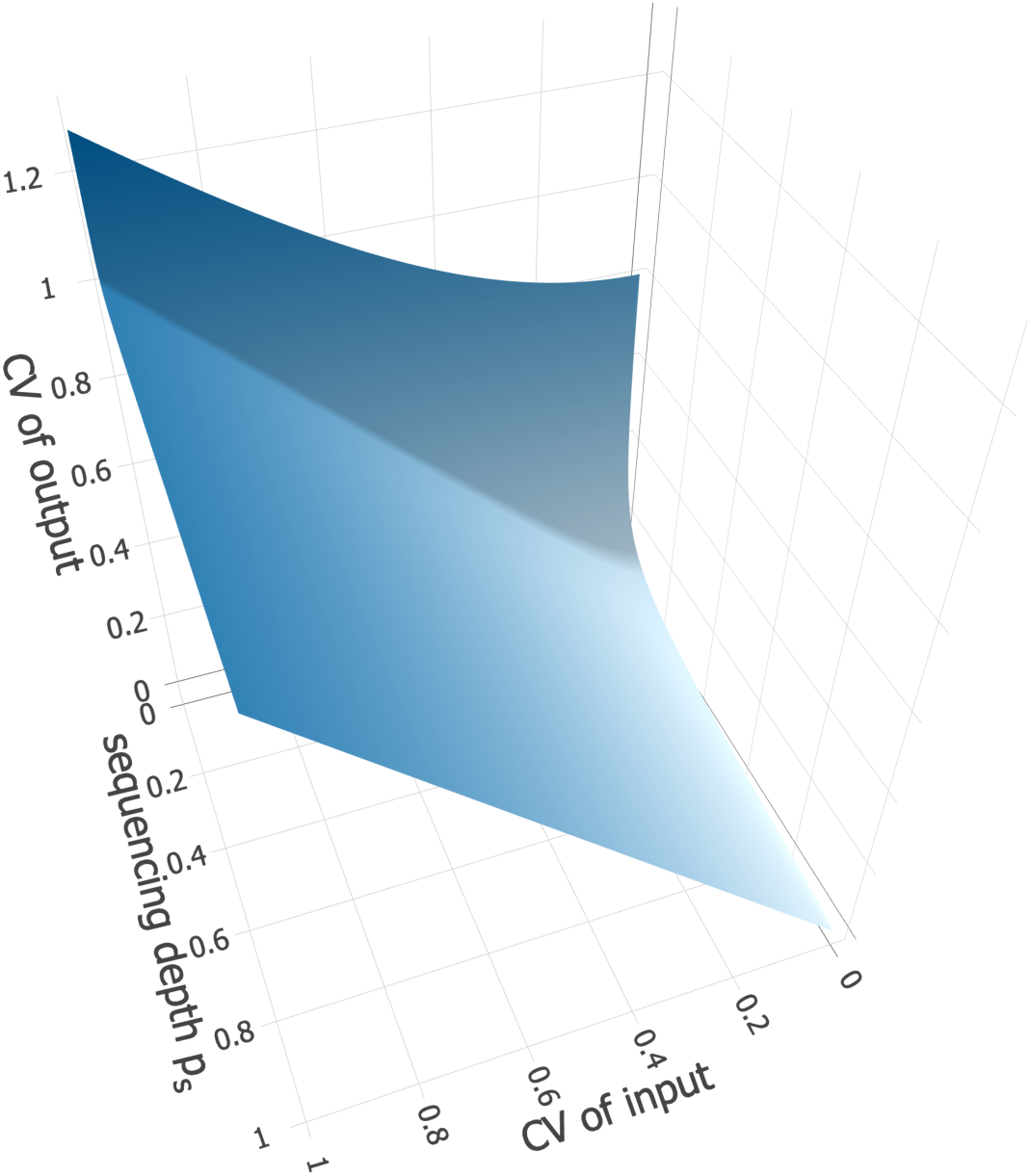
Effects of changes in input CV and success probability *p*_*s*_ on the output CV in three dimensions.

**Fig. S6.**
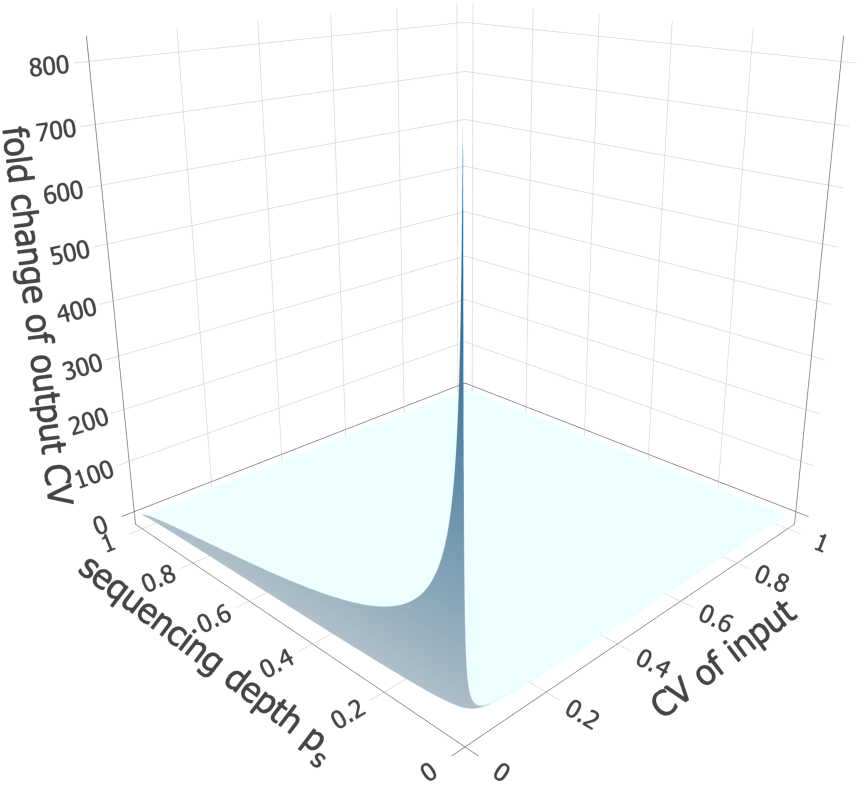
Effects of changes in input CV and success probability *p*_*s*_ on the fold change of the output CV compared to input CV in three dimensions.

